## Supplemental Data for "Binding stoichiometry and structural model of the HIV-1 Rev/Importin β complex"

### Supplementary Data

**Table S1. Summary of $IC_{50}$ values determined in FP competition assays[1].**

| Rev | WT | | R1 | | R2 | | R3 | | R4 | | R5 | |
|---|---|---|---|---|---|---|---|---|---|---|---|---|
| IC50 (μM) | Mean ± SD | N | Mean ± SD | N | Mean ± SD | N | Mean ± SD | N | Mean ± SD | N | Mean ± SD | N |
| **ImpB WT** | 0.502 ± 0.022 | 34 | 1.609 ± 0.052 | 6 | 4.971 ± 0.268 | 16 | 4.860 ± 0.325 | 22 | 4.852 ± 0.134 | 13 | 0.846 ± 0.023 | 6 |
| **B1** | 0.517 ± 0.041 | 6 | 1.399 ± 0.091 | 4 | 2.994 ± 0.250 | 6 | 2.873 ± 0.130 | 6 | 2.840 ± 0.126 | 6 | 0.549 ± 0.055 | 4 |
| **B2** | 0.549 ± 0.086 | 12 | 1.414 ± 0.061 | 4 | 1.649 ± 0.157 | 12 | 1.703 ± 0.208 | 11 | 1.533 ± 0.117 | 14 | 0.844 ± 0.095 | 4 |
| **B3** | 0.431 ± 0.050 | 12 | 1.481 ± 0.048 | 4 | 4.185 ± 0.496 | 10 | 3.298 ± 0.562 | 12 | 3.226 ± 0.213 | 6 | 0.670 ± 0.035 | 4 |
| **B4** | 0.441 ± 0.076 | 14 | 1.385 ± 0.022 | 4 | 4.188 ± 0.493 | 6 | 3.145 ± 0.446 | 14 | 3.080 ± 0.260 | 6 | 0.665 ± 0.011 | 4 |
| **B5** | 0.440 ± 0.021 | 6 | 1.295 ± 0.064 | 4 | 4.854 ± 0.509 | 6 | 4.807 ± 0.541 | 5 | 4.444 ± 0.304 | 6 | 0.844 ± 0.056 | 4 |
| **B6** | 0.433 ± 0.065 | 6 | 1.584 ± 0.174 | 4 | 4.538 ± 0.720 | 6 | 3.865 ± 0.430 | 6 | 3.960 ± 0.494 | 6 | 0.643 ± 0.070 | 4 |
| **B7** | 0.440 ± 0.082 | 6 | 1.575 ± 0.171 | 4 | 4.849 ± 0.849 | 6 | 4.727 ± 0.749 | 6 | 4.404 ± 0.655 | 6 | 0.866 ± 0.025 | 4 |

| Rev | WT | | R2 | | R35D | | R38D | | R39D | |
|---|---|---|---|---|---|---|---|---|---|---|
| IC50 (μM) | Mean ± SD | N | Mean ± SD | N | Mean ± SD | N | Mean ± SD | N | Mean ± SD | N |
| **ImpB WT** | 0.502 ± 0.022 | 34 | 4.971 ± 0.268 | 16 | 0.883 ± 0.028 | 10 | 0.802 ± 0.023 | 10 | 0.928 ± 0.048 | 10 |
| **B2** | 0.549 ± 0.086 | 12 | 1.649 ± 0.157 | 12 | 0.788 ± 0.146 | 6 | 0.771 ± 0.092 | 6 | 0.918 ± 0.107 | 6 |
| **D288R** | 0.629 ± 0.031 | 6 | 2.482 ± 0.161 | 6 | 0.926 ± 0.149 | 6 | 0.832 ± 0.125 | 5 | 0.998 ± 0.059 | 6 |
| **E289R** | 0.530 ± 0.070 | 6 | 4.139 ± 0.184 | 6 | 0.820 ± 0.076 | 6 | 0.744 ± 0.073 | 6 | 0.855 ± 0.074 | 6 |
| **D292R** | 0.531 ± 0.057 | 6 | 3.658 ± 0.109 | 6 | 0.752 ± 0.027 | 6 | 0.664 ± 0.036 | 6 | 0.794 ± 0.042 | 6 |
| **E299R** | 0.495 ± 0.067 | 6 | 4.487 ± 0.562 | 6 | 0.738 ± 0.029 | 6 | 0.722 ± 0.088 | 6 | 0.810 ± 0.059 | 6 |

| Rev | WT | | R3 | | R41D | | R44D | | R48D | |
|---|---|---|---|---|---|---|---|---|---|---|
| IC50 (μM) | Mean ± SD | N | Mean ± SD | N | Mean ± SD | N | Mean ± SD | N | Mean ± SD | N |
| **ImpB WT** | 0.502 ± 0.022 | 34 | 4.860 ± 0.325 | 22 | 0.828 ± 0.017 | 15 | 0.607 ± 0.021 | 14 | 0.551 ± 0.030 | 15 |
| **B2** | 0.549 ± 0.086 | 12 | 1.703 ± 0.208 | 11 | 0.604 ± 0.112 | 8 | 0.592 ± 0.037 | 4 | 0.541 ± 0.124 | 4 |
| **D288R** | 0.629 ± 0.031 | 6 | 2.398 ± 0.383 | 4 | 0.741 ± 0.189 | 8 | 0.740 ± 0.053 | 4 | 0.547 ± 0.125 | 4 |
| **E289R** | 0.530 ± 0.070 | 6 | 4.046 ± 0.979 | 4 | 0.678 ± 0.181 | 8 | 0.671 ± 0.066 | 4 | 0.495 ± 0.147 | 4 |
| **D292R** | 0.531 ± 0.057 | 6 | 3.804 ± 1.054 | 4 | 0.687 ± 0.205 | 8 | 0.688 ± 0.110 | 4 | 0.501 ± 0.157 | 4 |
| **E299R** | 0.495 ± 0.067 | 6 | 5.428 ± 1.153 | 4 | 0.831 ± 0.166 | 8 | 0.692 ± 0.064 | 4 | 0.495 ± 0.062 | 4 |
| **B3** | 0.431 ± 0.050 | 12 | 3.298 ± 0.562 | 12 | 0.691 ± 0.173 | 8 | 0.495 ± 0.074 | 6 | 0.329 ± 0.083 | 8 |
| **D339R** | 0.435 ± 0.095 | 6 | 3.125 ± 0.702 | 6 | 0.694 ± 0.146 | 8 | 0.554 ± 0.039 | 6 | 0.491 ± 0.081 | 8 |
| **D340R** | 0.537 ± 0.126 | 6 | 4.702 ± 0.982 | 6 | 0.860 ± 0.223 | 8 | 0.689 ± 0.133 | 6 | 0.550 ± 0.102 | 8 |
| **B4** | 0.441 ± 0.076 | 14 | 3.145 ± 0.446 | 14 | 0.700 ± 0.134 | 10 | 0.490 ± 0.116 | 8 | 0.316 ± 0.092 | 8 |
| **E437R** | 0.394 ± 0.079 | 8 | 3.753 ± 0.397 | 8 | 0.672 ± 0.166 | 10 | 0.493 ± 0.134 | 8 | 0.407 ± 0.122 | 8 |
| **E479R** | 0.509 ± 0.092 | 4 | 4.510 ± 1.587 | 8 | 0.888 ± 0.107 | 6 | 0.763 ± 0.092 | 4 | 0.585 ± 0.032 | 4 |
| **E534R** | 0.350 ± 0.018 | 5 | 4.311 ± 0.643 | 7 | 0.711 ± 0.147 | 9 | 0.581 ± 0.116 | 7 | 0.481 ± 0.127 | 8 |

| Rev | WT | | R4 | | R42D | | R43D | | R46D | |
|---|---|---|---|---|---|---|---|---|---|---|
| IC50 (μM) | Mean ± SD | N | Mean ± SD | N | Mean ± SD | N | Mean ± SD | N | Mean ± SD | N |
| **ImpB WT** | 0.502 ± 0.022 | 34 | 4.852 ± 0.134 | 13 | 0.646 ± 0.079 | 8 | 0.583 ± 0.027 | 8 | 0.579 ± 0.011 | 8 |
| **B2** | 0.549 ± 0.086 | 12 | 1.533 ± 0.117 | 14 | 0.481 ± 0.120 | 8 | 0.489 ± 0.052 | 8 | 0.440 ± 0.070 | 8 |
| **D288R** | 0.629 ± 0.031 | 6 | 2.065 ± 0.210 | 8 | 0.254 ± 0.026 | 6 | 0.387 ± 0.134 | 8 | 0.272 ± 0.023 | 6 |
| **E289R** | 0.530 ± 0.070 | 6 | 3.975 ± 0.169 | 8 | 0.539 ± 0.073 | 8 | 0.522 ± 0.047 | 8 | 0.452 ± 0.028 | 7 |
| **D292R** | 0.531 ± 0.057 | 6 | 3.907 ± 0.876 | 8 | 0.580 ± 0.143 | 8 | 0.515 ± 0.081 | 8 | 0.476 ± 0.105 | 8 |
| **E299R** | 0.495 ± 0.067 | 6 | 4.090 ± 0.218 | 7 | 0.494 ± 0.152 | 8 | 0.440 ± 0.120 | 8 | 0.417 ± 0.077 | 8 |

[1] Experimental values shown more than once are in normal font the first time they appear in the table and in grey upon subsequent instances.

### Table S2. Summary of initial HADDOCK docking experiments with Rev at the N- or C-site.

#### A. Docking parameters

**General parameters**

| | |
|---|---|
| PDB entries used | 2X7L (Rev), 1UKL (Imp$\beta$) |
| Active residues[(1)] | R42$^{Rev}$, R46$^{Rev}$, D288$^{Imp\beta}$ with (experiments 3-4) or without (experiments 1-2) E437$^{Imp\beta}$ |
| Passive residues[(1)] | All solvent-accessible residues (except those designated as active) on Rev helical hairpin (resi. 9-65) and concave inner surface of Imp$\beta$ [(2)] |

**Distance restraints**

| | Experiment 1 | Experiment 2 | Experiment 3 | Experiment 4 |
|---|---|---|---|---|
| Compensatory Mutagenesis | D288$^{Imp\beta}$(C$\gamma$) : R42$^{Rev}$(C$\zeta$) $\leq$ 5 Å<br>D288$^{Imp\beta}$(C$\gamma$) : R46$^{Rev}$(C$\zeta$) $\leq$ 5 Å | Same as Expt. 1 | D288$^{Imp\beta}$(C$\gamma$) : R42$^{Rev}$(C$\zeta$) $\leq$ 5 Å<br>D288$^{Imp\beta}$(C$\gamma$) : R46$^{Rev}$(C$\zeta$) $\leq$ 5 Å<br>E437$^{Imp\beta}$(C$\delta$) : R48$^{Rev}$(C$\zeta$) $\leq$ 6 Å | Same as Expt. 3 |
| BS3 Crosslinking | K23(C$\beta$)$^{Imp\beta}$ : K20(C$\beta$)$^{Rev}$ $\leq$ 30 Å<br>K62(C$\beta$)$^{Imp\beta}$ : K20(C$\beta$)$^{Rev}$ $\leq$ 30 Å<br>K68(C$\beta$)$^{Imp\beta}$ : K20(C$\beta$)$^{Rev}$ $\leq$ 30 Å | K854(C$\beta$)$^{Imp\beta}$ : K20(C$\beta$)$^{Rev}$ $\leq$ 30 Å<br>K857(C$\beta$)$^{Imp\beta}$ : K20(C$\beta$)$^{Rev}$ $\leq$ 30 Å<br>K859(C$\beta$)$^{Imp\beta}$ : K20(C$\beta$)$^{Rev}$ $\leq$ 30 Å<br>K867(C$\beta$)$^{Imp\beta}$ : K20(C$\beta$)$^{Rev}$ $\leq$ 30 Å<br>K873(C$\beta$)$^{Imp\beta}$ : K20(C$\beta$)$^{Rev}$ $\leq$ 30 Å | Same as Expt. 1 | Same as Expt. 2 |

#### B. Docking results

| | Experiment 1 | Experiment 2 | Experiment 3 | Experiment 4 |
|---|---|---|---|---|
| Total structures clustered: [(3)] | 143 | 177 | 84 | 189 |
| No. clusters: | 9 | 11 | 7 | 7 |

**Docking Statistics [(4)]:**

| | Rank | Cluster ID [(5)] | Cluster size | Haddock Score (kcal/mol) | Z-score | rmsd from LES (Å) [(6)] | Van der Waals Energy (kcal/mol) | Electrostatic Energy (kcal/mol) | Desolvation Energy (kcal/mol) | Restraints Violation Energy (kcal/mol) | Buried Surface Area (Å$^2$) |
|---|---|---|---|---|---|---|---|---|---|---|---|
| **Expt.1** | 1 | 1 | 61 | -160.2 ± 3.6 | -1.5 | 1.2 ± 0.2 | -37.9 ± 1.5 | -775.9 ± 71.5 | 32.8 ± 15.8 | 0.3 ± 0.3 | 1864.9 ± 110.6 |
| | 2 | 3 | 15 | -149.7 ± 14.2 | -1.1 | 0.9 ± 0.5 | -42.7 ± 7.0 | -723.4 ± 63.5 | 37.7 ± 7.5 | 0.0 ± 0.1 | 1876.8 ± 176.1 |
| | 3 | 4 | 9 | -145.5 ± 15.0 | -1.0 | 1.6 ± 0.3 | -37.1 ± 7.8 | -672.8 ± 67.8 | 26.2 ± 8.1 | 0.0 ± 0.0 | 2121.8 ± 173.4 |
| | 4 | 9 | 4 | -123.8 ± 27.7 | 0.1 | 2.1 ± 0.1 | -33.4 ± 10.7 | -642.4 ± 102.0 | 38.1 ± 8.6 | 0.0 ± 0.0 | 2058.5 ± 268.9 |
| | 5 | 2 | 34 | -122.9 ± 16.0 | 0.1 | 2.0 ± 0.4 | -26.1 ± 9.2 | -516.9 ± 72.5 | 6.6 ± 6.5 | 0.0 ± 0.0 | 1360.0 ± 259.5 |
| | 6 | 5 | 7 | -106.4 ± 8.9 | 0.6 | 8.0 ± 0.1 | -35.6 ± 9.5 | -425.7 ± 63.9 | 11.4 ± 8.3 | 29.0 ± 17.0 | 1728.4 ± 56.0 |
| | 7 | 8 | 4 | -100.8 ± 13.5 | 0.8 | 2.0 ± 0.2 | -32.5 ± 3.6 | -428.5 ± 11.1 | 17.4 ± 10.5 | 0.1 ± 0.1 | 1620.8 ± 169.2 |
| | 8 | 6 | 5 | -99.3 ± 12.2 | 0.8 | 1.8 ± 0.2 | -25.9 ± 9.8 | -485.1 ± 55.2 | 23.6 ± 3.0 | 0.0 ± 0.0 | 1674.0 ± 247.3 |
| | 9 | 7 | 4 | -77.5 ± 16.1 | 1.7 | 1.7 ± 0.2 | -13.8 ± 3.4 | -395.2 ± 53.7 | 15.3 ± 9.5 | 0.0 ± 0.0 | 1018.2 ± 102.2 |
| **Expt.2** | 1 | 1 | 68 | -169.1 ± 11.7 | -2.3 | 0.7 ± 0.4 | -48.7 ± 5.7 | -584.8 ± 102.0 | 3.5 ± 8.2 | 0.1 ± 0.1 | 2109.8 ± 115.6 |
| | 2 | 8 | 6 | -139.2 ± 18.1 | -0.8 | 3.4 ± 0.1 | -42.3 ± 5.4 | -643.4 ± 45.4 | 28.8 ± 11.5 | 29.2 ± 16.0 | 2042.9 ± 95.2 |
| | 3 | 2 | 48 | -136.0 ± 1.9 | -0.6 | 1.9 ± 0.2 | -39.9 ± 8.2 | -494.2 ± 53.1 | 2.7 ± 8.9 | 0.0 ± 0.0 | 1718.7 ± 69.1 |
| | 4 | 3 | 11 | -126.2 ± 3.3 | -0.1 | 2.4 ± 0.1 | -38.5 ± 5.0 | -477.1 ± 94.4 | -0.7 ± 19.1 | 83.5 ± 21.1 | 1821.9 ± 154.8 |
| | 5 | 6 | 7 | -124.0 ± 2.7 | 0.0 | 3.2 ± 0.1 | -27.1 ± 6.2 | -552.3 ± 51.5 | 12.8 ± 8.1 | 7.8 ± 12.6 | 1518.3 ± 101.6 |
| | 6 | 4 | 9 | -116.6 ± 7.4 | 0.4 | 2.6 ± 0.4 | -34.6 ± 5.3 | -469.8 ± 61.5 | 11.9 ± 16.6 | 0.0 ± 0.0 | 1572.2 ± 172.6 |
| | 7 | 7 | 6 | -111.7 ± 10.6 | 0.6 | 4.8 ± 0.1 | -43.3 ± 8.4 | -511.8 ± 26.6 | 11.9 ± 16.6 | 173.2 ± 6.3 | 1951.9 ± 63.9 |
| | 8 | 5 | 9 | -107.0 ± 18.5 | 0.9 | 1.7 ± 0.1 | -24.6 ± 4.2 | -346.4 ± 95.5 | -13.1 ± 6.8 | 0.5 ± 0.5 | 1198.4 ± 73.8 |
| | 9 | 9 | 5 | -105.2 ± 16.5 | 1.0 | 2.1 ± 0.3 | -32.1 ± 10.9 | -368.7 ± 77.5 | -2.0 ± 10.9 | 26.4 ± 32.9 | 1548.7 ± 191.1 |
| | 10 | 10 | 4 | -102.2 ± 30.6 | 1.1 | 2.9 ± 0.1 | -45.4 ± 4.8 | -356.8 ± 74.3 | 13.4 ± 15.2 | 11.8 ± 19.9 | 1692.1 ± 171.5 |
| | 11 | 11 | 4 | -98.0 ± 23.5 | 1.3 | 2.0 ± 0.1 | -44.2 ± 10.2 | -395.8 ± 89.1 | 25.3 ± 9.6 | 0.0 ± 0.1 | 1696.3 ± 250.8 |
| **Expt. 3** | 1 | 1 | 33 | -142.4 ± 15.6 | -1.7 | 0.7 ± 0.4 | -62.1 ± 8.8 | -627.7 ± 90.7 | 8.9 ± 13.9 | 363.5 ± 28.3 | 2479.9 ± 117.0 |
| | 2 | 2 | 17 | -113.7 ± 4.7 | -0.8 | 8.7 ± 0.2 | -34.1 ± 4.7 | -582.0 ± 39.8 | 21.8 ± 5.7 | 150.2 ± 3.0 | 1932.6 ± 59.9 |
| | 3 | 3 | 13 | -109.2 ± 18.8 | -0.6 | 1.3 ± 0.1 | -42.6 ± 3.9 | -622.1 ± 123.8 | 23.1 ± 11.2 | 347.2 ± 51.2 | 2284.3 ± 97.8 |
| | 4 | 5 | 5 | -77.6 ± 12.6 | 0.5 | 6.9 ± 0.0 | -34.6 ± 8.4 | -482.4 ± 68.8 | 24.5 ± 24.1 | 290.1 ± 16.1 | 1783.7 ± 179.7 |
| | 5 | 4 | 8 | -73.5 ± 11.0 | 0.6 | 9.8 ± 0.2 | -35.0 ± 5.8 | -543.6 ± 42.1 | 16.6 ± 8.2 | 536.8 ± 40.2 | 1568.3 ± 114.1 |
| | 6 | 6 | 4 | -71.4 ± 16.5 | 0.7 | 2.9 ± 0.1 | -47.2 ± 5.7 | -526.5 ± 28.7 | 56.7 ± 8.6 | 243.6 ± 20.5 | 2312.0 ± 84.6 |
| | 7 | 7 | 4 | -49.2 ± 14.7 | 1.4 | 2.6 ± 0.2 | -23.4 ± 10.4 | -523.6 ± 89.9 | 30.1 ± 9.2 | 488.8 ± 20.6 | 1613.9 ± 198.2 |
| **Expt. 4** | 1 | 1 | 73 | -152.7 ± 9.4 | -1.9 | 2.1 ± 0.1 | -51.1 ± 3.1 | -546.2 ± 61.3 | -0.7 ± 16.2 | 82.9 ± 18.1 | 2037.3 ± 76.3 |
| | 2 | 5 | 13 | -135.5 ± 8.7 | -0.3 | 1.7 ± 0.1 | -29.9 ± 1.8 | -652.4 ± 66.4 | 13.7 ± 6.6 | 112.1 ± 16.2 | 1750.2 ± 162.6 |
| | 3 | 2 | 43 | -134.2 ± 7.6 | -0.2 | 1.6 ± 0.3 | -43.3 ± 1.8 | -518.9 ± 52.7 | 11.8 ± 10.5 | 10.7 ± 16.0 | 1776.9 ± 102.0 |
| | 4 | 3 | 27 | -133.8 ± 6.6 | -0.1 | 2.2 ± 0.1 | -34.2 ± 1.6 | -642.2 ± 78.0 | 226.1 ± 13.8 | 27.1 ± 14.7 | 2004.5 ± 97.4 |
| | 5 | 6 | 10 | -132.3 ± 23.8 | 0.0 | 0.9 ± 0.6 | -37.9 ± 1.2 | -549.2 ± 82.5 | 15.4 ± 15.7 | 1.6 ± 2.0 | 1701.2 ± 102.4 |
| | 6 | 7 | 4 | -121.4 ± 16.0 | 1.1 | 1.5 ± 0.1 | -32.7 ± 8.6 | -559.3 ± 88.4 | 19.7 ± 12.7 | 34.1 ± 39.4 | 1710.7 ± 212.3 |
| | 7 | 4 | 19 | -118.0 ± 4.2 | 1.4 | 1.8 ± 0.2 | -40.5 ± 4.3 | -494.2 ± 55.3 | 19.4 ± 12.3 | 18.8 ± 31.0 | 1739.5 ± 122.0 |

[1] Active residues are those designated as being explicitly located in the intermolecular interface, while passive residues are those designated as potentially but not necessarily in the interface.

[2] These consist of Imp$\beta$ residues 8-19, 21-23, 25- 27, 29-31, 33-37, 50-52, 55-56, 59-60, 62-64, 66-79, 81-82, 102-105, 107-108, 110-111, 114-115, 118-119, 122-125, 140-145, 147-149, 152, 155-156, 158-165, 167-169, 184, 186-189, 192-193, 195-197, 199-200, 203-204, 206-207, 209- 212, 229-232, 235-236, 239, 242-243, 246-251, 253-254, 271-275, 277-278, 280-281, 284-285, 288-293, 295-299, 301-313, 315-317, 328-344, 346-347, 350-351, 353-354, 357-362, 377-385, 388, 392, 395-400, 402-403, 420-423, 425-427, 429-430, 434, 436-441, 444, 460, 463-466, 468-469, 471-472, 475-477, 479-484, 486-487, 489-494, 517-518, 520-523, 525-527, 530-531, 533-534, 537-538, 541, 544, 565-572, 574-576, 578-579, 581-583, 585-586, 589-590, 593-598, 613, 616-627, 630, 633-634, 636-642, 644, 660-666, 668-669, 671-673, 675-676, 678-684, 702-709, 711-712, 715-716, 718-719, 722-727, 748-757, 759-760, 763-764, 767-768, 770-771, 774-784, 786, 805-806, 808-813, 816-817, 819-821, 824, 827-828, 829-831, 834-835, 837-839, 841-842, 844-845, 848-849, 851-857, 859-861, 863-864, 866-876.

[3] Number of structures clustered (out of 200) in the final refinement stage of docking. Structures are clustered if they have a pairwise RMSD < 7.5 Å.

[4] Docking statistics reported (HADDOCK score, rmsd, energy terms and buried surface area) represent the mean value and standard deviation for the four lowest energy structures in each cluster.

[5] Cluster ID is the rank of the cluster when ranked according to cluster size.

[6] LES: lowest-energy structure.

### Table S3. Statistics for rigid body docking of Rev at the C-site.

**A. Cβ-Cβ Distance constraints**

| Compensatory Mutagenesis | BS3 Crosslinking |
|---|---|
| $D288^{Imp\beta}$ : $R42^{Rev} \leq 15$ Å | $K537^{Imp\beta}$ : $K20^{Rev} \leq 30$ Å |
| $D288^{Imp\beta}$ : $R43^{Rev} \leq 15$ Å | $K854^{Imp\beta}$ : $K20^{Rev} \leq 30$ Å |
| $D288^{Imp\beta}$ : $R46^{Rev} \leq 15$ Å | $K857^{Imp\beta}$ : $K20^{Rev} \leq 30$ Å |
| $E289^{Imp\beta}$ : $R46^{Rev} \leq 15$ Å | $K859^{Imp\beta}$ : $K20^{Rev} \leq 30$ Å |
| $E299^{Imp\beta}$ : $R42^{Rev} \leq 15$ Å | $K867^{Imp\beta}$ : $K20^{Rev} \leq 30$ Å |
| $E299^{Imp\beta}$ : $R43^{Rev} \leq 15$ Å | $K873^{Imp\beta}$ : $K20^{Rev} \leq 30$ Å |
| $E299^{Imp\beta}$ : $R46^{Rev} \leq 15$ Å | |
| $E437^{Imp\beta}$ : $R48^{Rev} \leq 15$ Å | |

**B. Docking Results**

| | Minimal distance between Impβ and Rev side chains (Å) (1) | | | | | | | | mean sc/sc | Change in Rev vs. Rank1(3) | |
|---|---|---|---|---|---|---|---|---|---|---|---|
| Rank | $D288^{Imp\beta}$ : $R42^{Rev}$ | $D288^{Imp\beta}$ : $R43^{Rev}$ | $D288^{Imp\beta}$ : $R46^{Rev}$ | $E289^{Imp\beta}$ : $R46^{Rev}$ | $E299^{Imp\beta}$ : $R42^{Rev}$ | $E299^{Imp\beta}$ : $R43^{Rev}$ | $E299^{Imp\beta}$ : $R46^{Rev}$ | $E437^{Imp\beta}$ : $R48^{Rev}$ | distance(2) (Å) | Angle (°) | Shift(4) (Å) |
| 1 | 2.96 | 2.30 | 3.55 | 8.01 | 5.61 | 12.77 | 5.47 | 3.24 | 4.61 | - | - |
| 2 | 2.30 | 2.30 | 2.72 | 6.11 | 7.14 | 12.93 | 4.72 | 8.83 | 4.66 | 60.7 | 2.9 |
| 3 | 2.69 | 2.30 | 2.30 | 7.23 | 9.13 | 14.61 | 7.42 | 3.98 | 4.87 | 15.0 | 4.4 |
| 4 | 3.77 | 2.30 | 5.13 | 7.57 | 3.48 | 13.44 | 2.83 | 7.06 | 5.10 | 33.0 | 3.5 |
| 5 | 4.71 | 2.30 | 5.17 | 9.56 | 6.43 | 8.56 | 3.55 | 4.16 | 5.15 | 33.0 | 1.9 |
| 6 | 4.79 | 3.37 | 2.55 | 7.86 | 8.00 | 14.27 | 6.05 | 3.52 | 5.37 | 29.0 | 4.7 |
| 7 | 5.03 | 2.30 | 6.68 | 9.63 | 2.30 | 11.12 | 2.39 | 4.94 | 5.41 | 0.0 | 5.7 |
| 8 | 4.54 | 2.60 | 6.33 | 10.06 | 2.64 | 12.14 | 3.11 | 9.06 | 5.80 | 29.0 | 5.0 |
| 9 | 7.47 | 2.30 | 6.81 | 8.79 | 2.30 | 8.32 | 2.30 | 6.98 | 5.97 | 33.0 | 5.3 |
| 10 | 7.84 | 2.30 | 6.00 | 7.98 | 5.28 | 7.24 | 2.30 | 7.19 | 6.00 | 29.0 | 3.0 |
| 11 | 3.72 | 3.93 | 4.80 | 9.86 | 9.34 | 13.61 | 7.37 | 5.35 | 6.10 | 30.0 | 7.3 |
| 12 | 5.47 | 2.30 | 6.61 | 10.66 | 9.41 | 9.06 | 4.09 | 7.44 | 6.34 | 66.8 | 6.4 |
| 13 | 5.47 | 3.71 | 6.57 | 11.34 | 7.52 | 11.05 | 4.58 | 6.00 | 6.48 | 44.0 | 6.6 |
| 14 | 6.70 | 4.51 | 6.54 | 10.75 | 2.67 | 12.82 | 3.01 | 8.84 | 6.72 | 32.1 | 3.7 |
| 15 | 8.92 | 2.30 | 5.81 | 8.93 | 11.55 | 9.84 | 5.66 | 6.36 | 7.24 | 33.0 | 7.4 |
| | | | | | | | | | **Mean:** | **33.4 ± 15.8** | **4.8 ± 1.7** |

[1] Rotamer combinations yielding a side chain separation less than 2.3 Å were assigned a minimal distance of 2.3 Å.
[2] $\Delta\Delta p IC_{50}$-weighted mean distance between side chains for the 8 pairs of Impβ and Rev residues, calculated as $\Sigma(\Delta\Delta p IC_{50,i} * D_i)/\Sigma(\Delta\Delta p IC_{50,i})$, where $D_i$ is the minimal distance for the $i^{th}$ pair of Impβ and Rev side chains and $\Delta\Delta p IC_{50,i}$ is the corresponding $\Delta\Delta p IC_{50}$ value.
[3] The change in the orientation and position of Rev compared to those in the top-ranked structure.
[4] Distance between the centroids of the two Rev monomers compared.

### Table S4. Summary of final HADDOCK docking experiment with Rev at the C-site.

#### A. Docking parameters

**General parameters**

| | |
|---|---|
| PDB entries used | 2X7L (Rev), 1UKL (Impβ) |
| Active residues[1] | Rev residues R42, R43, R46, R48; Impβ residues D288, D289, D299, E437 |
| Passive residues[1] | All solvent-accessible residues (except those designated as active) on Rev helical hairpin (resi. 9-65) and concave inner surface of Impβ [2] |

**Distance restraints**

| | **Compensatory Mutagenesis** | **BS3 Crosslinking** |
|---|---|---|
| | D288$^{Imp\beta}$(Cγ) : R42$^{Rev}$(Cζ) ≤ 5 Å | K537(Cβ)$^{Imp\beta}$ : K20(Cβ)$^{Rev}$ ≤ 30 Å |
| | D288$^{Imp\beta}$(Cγ) : R43$^{Rev}$(Cζ) ≤ 5 Å | K854(Cβ)$^{Imp\beta}$ : K20(Cβ)$^{Rev}$ ≤ 30 Å |
| | D288$^{Imp\beta}$(Cγ) : R46$^{Rev}$(Cζ) ≤ 5 Å | K857(Cβ)$^{Imp\beta}$ : K20(Cβ)$^{Rev}$ ≤ 30 Å |
| | E289$^{Imp\beta}$(Cδ) : R46$^{Rev}$(Cζ) ≤ 8 Å | K859(Cβ)$^{Imp\beta}$ : K20(Cβ)$^{Rev}$ ≤ 30 Å |
| | E299$^{Imp\beta}$(Cδ) : R42$^{Rev}$(Cζ) ≤ 8 Å | K867(Cβ)$^{Imp\beta}$ : K20(Cβ)$^{Rev}$ ≤ 30 Å |
| | E299$^{Imp\beta}$(Cδ) : R43$^{Rev}$(Cζ) ≤ 8 Å | K873(Cβ)$^{Imp\beta}$ : K20(Cβ)$^{Rev}$ ≤ 30 Å |
| | E299$^{Imp\beta}$(Cδ) : R46$^{Rev}$(Cζ) ≤ 6 Å | |
| | E437$^{Imp\beta}$(Cδ) : R48$^{Rev}$(Cζ) ≤ 6 Å | |

#### B. Docking results [3]

| **Rank** | **Cluster ID** [4] | **Cluster Size** [5] | **Haddock Score (kcal/mol)** | **Z-score** | **rmsd vs. LES [6] (Å)** | **Van der Waals Energy (kcal/mol)** | **Electrostatic Energy (kcal/mol)** | **Desolvation Energy (kcal/mol)** | **Restraints Violation Energy (kcal/mol)** | **Buried Surface Area (Å²)** | **Change in Rev vs. LES$_1$ [7]** Angle (°) | Shift[8] (Å) |
|---|---|---|---|---|---|---|---|---|---|---|---|---|
| 1 | 1 | 112 | -166.7 ± 4.6 | -1.9 | 0.7 ± 0.4 | -36.8 ± 3.7 | -781.9 ± 50.9 | 10.2 ± 6.3 | 162.9 ± 39.6 | 1927.1 ± 76.7 | 7.2 ± 2.5 | 1.5 ± 0.5 |
| 2 | 5 | 9 | -146.2 ± 11.8 | -1.0 | 1.3 ± 0.1 | -41.2 ± 5.1 | -825.8 ± 41.2 | 40.7 ± 15.7 | 194.1 ± 17.8 | 2326.1 ± 64.0 | 46.5 ± 7.5 | 1.8 ± 0.8 |
| 3 | 3 | 19 | -123.7 ± 6.9 | 0.0 | 1.3 ± 0.3 | -29.1 ± 9.5 | -681.8 ± 88.2 | 20.3 ± 21.2 | 214.6 ± 57.5 | 1610.3 ± 103.0 | 24.2 ± 5.2 | 3.4 ± 1.7 |
| 4 | 2 | 27 | -116.7 ± 4.5 | 0.3 | 1.7 ± 0.4 | -31.1 ± 6.5 | -624.9 ± 88.9 | 23.0 ± 12.7 | 164.2 ± 27.5 | 1635.2 ± 138.9 | 58.2 ± 5.3 | 4.6 ± 0.5 |
| 5 | 4 | 16 | -112.2 ± 4.4 | 0.5 | 2.0 ± 0.2 | -20.6 ± 3.5 | -632.1 ± 52.1 | 17.4 ± 15.3 | 174.9 ± 52.3 | 1484.8 ± 163.1 | 34.9 ± 0.9 | 3.0 ± 0.3 |
| 6 | 6 | 6 | -106.9 ± 7.2 | 0.7 | 2.1 ± 0.4 | -46.3 ± 7.1 | -519.6 ± 50.4 | 17.1 ± 18.3 | 261.9 ± 27.9 | 1726.7 ± 148.6 | 24.3 ± 9.3 | 5.8 ± 1.0 |
| 7 | 7 | 5 | -94.3 ± 21.2 | 1.3 | 1.9 ± 0.1 | -40.7 ± 6.3 | -541.3 ± 65.2 | 25.6 ± 10.9 | 290.5 ± 45.1 | 2068.5 ± 97.6 | 16.9 ± 1.4 | 5.6 ± 0.2 |
| | **Total:** | **194** | | | | | | | | **Mean:** | **34.2 ± 16.2** | **4.0 ± 1.6** |

[1] Active residues as those designated as being explicitly located in the intermolecular interface, while passive residues are those designated as potentially but not necessarily in the interface.

[2] See footnote (2) of Table S2 for explicit list of residues.

[3] Docking statistics reported (HADDOCK score, rmsd, energy terms and buried surface area) represent the mean value and standard deviation for the four lowest energy structures in each cluster.

[4] Cluster ID is the rank of the cluster when ranked according to cluster size.

[5] Number of structures clustered (out of a total of 200 tested) in the final refinement stage of docking. Structures are clustered if they have a pairwise RMSD < 7.5 Å.

[6] LES: lowest-energy structure.

[7] The change in the orientation and position of Rev in the four lowest-energy structures of each cluster compared to those in the lowest-energy structure of cluster 1.

[8] Distance between the centroids of the two Rev monomers compared.

**Table S5. Quality control of Impβ and Rev constructs by LC/ESI mass spectrometric analysis.**

| Protein | Mutations | Expected average mass (Da) | Observed average mass (Da) | Δ mass (Observed - Expected) (Da) |
|---|---|---|---|---|
| Impβ (WT) | – | 97,298.6 | 97,299.6 | 1.0 |
| **His-Impβ** | | | | |
| WT | – | 100,296.9 | 100,299.2 | 2.3 |
| B1 | E152R E203R E274R E281R | 100,405.2 | 100,403.3 | -1.9 |
| B2 | D288R E289R D292R E299R | 100,433.3 | 100,432.4 | -0.9 |
| B3 | D339R D340R | 100,379.1 | 100,373.4 | -5.7 |
| B4 | E437R E479R E534R | 100,378.1 | 100,380.3 | 2.2 |
| B5 | E483R D486R D490R | 100,405.9 | 100,404.2 | -1.7 |
| B6 | E530R D579R E626R | 100,390.1 | 100,392.2 | 2.1 |
| B8 | D753R D756R E760R | 100,405.9 | 100,404.2 | -1.7 |
| D288R | D288R | 100,338.0 | 100,342.6 | 4.6 |
| E289R | E289R | 100,324.0 | 100,328.3 | 4.3 |
| D292R | D292R | 100,338.0 | 100,343.0 | 5.0 |
| E299R | E299R | 100,324.0 | 100,327.5 | 3.5 |
| D339R | D339R | 100,338.0 | 100,340.2 | 2.2 |
| D340R | D340R | 100,338.0 | 100,340.9 | 2.9 |
| E437R | E437R | 100,324.0 | 100,331.7 | 7.7 |
| E479R | E479R | 100,324.0 | 100,331.2 | 7.2 |
| E534R | E534R | 100,324.0 | 100,331.7 | 7.7 |
| **Rev** | | | | |
| WT | – | 13,175.9 | 13,176.9 | 1.0 |
| OD* | V16D I55N | 13,263.8 | 13,263.9 | 0.1 |
| ODΔ | V16D I55N | 8,030.9 | 8,031.0 | 0.1 |
| R1 | R14D R17D K20D | 13,080.6 | 13,080.4 | -0.2 |
| R2 | R35D R38D R39D | 13 052.6 | 13,050.6 | -2.0 |
| R3 | R41D R44D R48D | 13,052.6 | 13,052.4 | -0.2 |
| R4 | R42D R43D R46D | 13,052.6 | 13,052.1 | -0.5 |
| R5 | R50D R58D | 13,093.7 | 13,093.3 | -0.4 |
| R35D | R35D | 13,134.8 | 13,134.9 | 0.1 |
| R38D | R38D | 13,134.8 | 13,134.1 | -0.7 |
| R39D | R39D | 13,134.8 | 13,134.0 | -0.8 |
| R41D | R41D | 13,134.8 | 13,135.0 | 0.2 |
| R42D | R42D | 13,134.8 | 13,135.1 | 0.3 |
| R43D | R43D | 13,134.8 | 13,134.8 | 0.0 |
| R44D | R44D | 13,134.8 | 13,135.1 | 0.3 |
| R46D | R46D | 13,134.8 | 13,134.8 | 0.0 |
| R48D | R48D | 13,134.8 | 13,135.1 | 0.3 |

* The $Rev^{OD}$ construct contains an additional Ala residue after the TEV cleavage site.

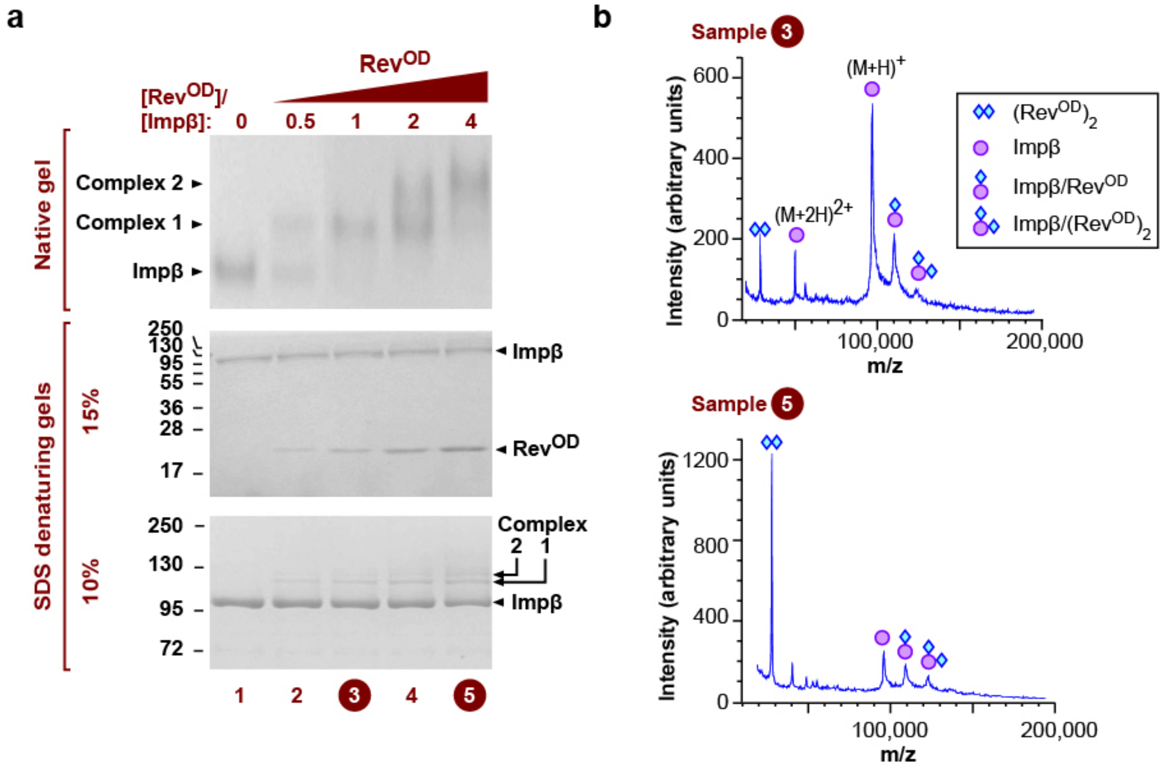


**Figure S1. Glutaraldehyde crosslinking and MALDI-TOF mass spectrometry indicate that Impβ binds two Rev monomers. a.** Impβ was incubated with Rev$^{OD}$ in the presence of glutaraldehyde and the mixture was analysed by native (top panel) and denaturing (bottom two panels) gel electrophoresis followed by Coomassie staining. The upper denaturing gel (containing 15% acrylamide:bis-acrylamide in a 37.5:1 ratio) allowed Impβ and Rev to be visualized on the same gel, while the lower gel (containing 10% acrylamide:bis-acrylamide) was used to resolve crosslinked species migrating close to Impβ. At Rev$^{OD}$ concentrations yielding a single gel shift by native gel electrophoresis, SDS-PAGE analysis revealed an additional band that migrated just above Impβ (bottom gel, compare lanes 1 and 3). At higher Rev$^{OD}$ concentrations that yielded a supershift on the native gel, a second more slowly migrating band was detected by SDS-PAGE (bottom gel, lanes 4-5). **b.** MALDI-TOF mass spectrometry analysis of samples corresponding to lanes 3 (upper profile) and 5 (lower profile) from panel **a** was performed. MALDI MS spectra showed peaks consistent with the molecular mass of free Impβ (indicated by a circle), Impβ crosslinked to either one or two Rev$^{OD}$ monomers (indicated by a circle attached to one or two diamonds, respectively), or crosslinked Rev$^{OD}$ dimers (indicated by two diamonds.)

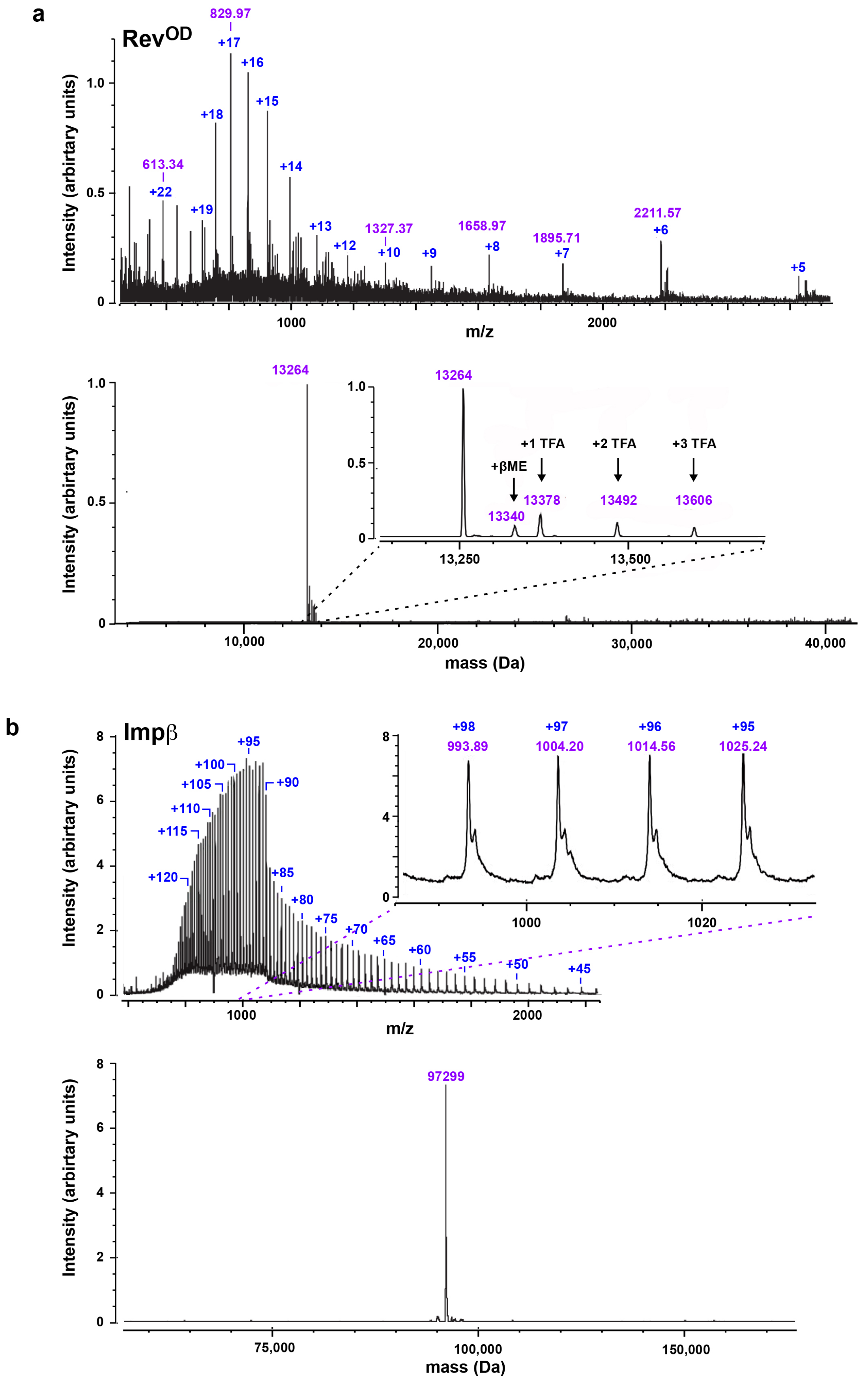


**Figure S2. LC/ESI-TOF MS analysis of (a) Rev$^{OD}$ and (b) Impβ.** The upper mass spectrum in each panel represents the raw ESI data while the lower panel shows the deconvoluted ESI spectrum. βME, β-mercaptoethanol. TFA, trifluoroacetic acid.

**a**

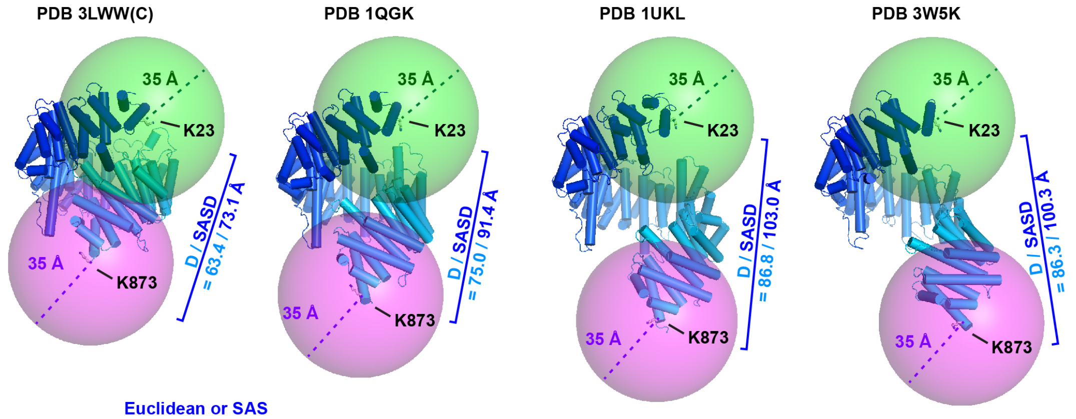


**b**

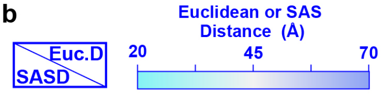


PDB 3LWW(C)

| | K23 | K62 | K68 | K206 | K537 | K854 | K857 | K859 | K867 | K873 |
|---|---|---|---|---|---|---|---|---|---|---|
| K23 | -- | 17.8 | 21.1 | 46.9 | 52.3 | 52.3 | 52.5 | 54.2 | 59.0 | 63.4 |
| K62 | 26.4 | -- | 11.1 | 30.0 | 49.3 | 42.1 | 44.9 | 45.3 | 50.9 | 55.3 |
| K68 | 21.1 | 11.1 | -- | 28.0 | 39.2 | 32.4 | 34.3 | 35.1 | 41.3 | 46.9 |
| K206 | 59.6 | 38.8 | 35.1 | -- | 51.1 | 29.9 | 36.7 | 33.9 | 38.9 | 42.3 |
| K537 | 56.4 | 52.3 | 43.6 | 51.1 | -- | 26.5 | 23.8 | 28.7 | 38.5 | 49.1 |
| K854 | 57.8 | 45.5 | 37.0 | 29.9 | 27.4 | -- | 8.9 | 7.9 | 19.6 | 29.5 |
| K857 | 62.9 | 47.6 | 38.8 | 38.8 | 23.8 | 17.0 | -- | 5.2 | 15.4 | 26.1 |
| K859 | 74.0 | 58.4 | 49.6 | 44.2 | 28.7 | 19.5 | 5.2 | -- | 12.2 | 22.6 |
| K867 | 84.1 | 70.5 | 61.8 | 44.1 | 40.2 | 25.3 | 18.9 | 18.6 | -- | 10.9 |
| K873 | 73.1 | 67.9 | 63.8 | 42.5 | 52.5 | 36.3 | 31.1 | 30.8 | 10.9 | -- |

PDB 1UKL

| | K23 | K62 | K68 | K206 | K537 | K854 | K857 | K859 | K867 | K873 |
|---|---|---|---|---|---|---|---|---|---|---|
| K23 | -- | 17.2 | 21.8 | 46.5 | 59.8 | 68.2 | 67.6 | 71.5 | 81.4 | 86.8 |
| K62 | 28.6 | -- | 11.7 | 30.1 | 57.0 | 59.8 | 61.4 | 64.3 | 74.8 | 80.3 |
| K68 | 21.8 | 11.7 | -- | 26.7 | 45.6 | 48.7 | 50.0 | 53.1 | 63.8 | 69.7 |
| K206 | 52.3 | 36.3 | 27.1 | -- | 53.4 | 44.0 | 49.7 | 50.0 | 60.6 | 65.6 |
| K537 | 63.9 | 60.6 | 48.8 | 56.2 | -- | 27.9 | 25.1 | 29.5 | 39.1 | 48.3 |
| K854 | 69.6 | 60.2 | 48.8 | 44.0 | 27.9 | -- | 9.1 | 7.2 | 19.3 | 27.5 |
| K857 | 67.6 | 64.5 | 53.1 | 51.4 | 25.1 | 13.4 | -- | 5.5 | 15.7 | 24.4 |
| K859 | 76.9 | 70.0 | 58.6 | 51.9 | 29.5 | 7.2 | 9.7 | -- | 12.6 | 21.3 |
| K867 | 91.4 | 84.5 | 73.1 | 65.3 | 39.3 | 19.3 | 20.8 | 13.9 | -- | 9.6 |
| K873 | 103.0 | 92.7 | 84.3 | 72.2 | 51.1 | 30.4 | 34.6 | 23.9 | 11.2 | -- |

PDB 1QGK

| | K23 | K62 | K68 | K206 | K537 | K854 | K857 | K859 | K867 | K873 |
|---|---|---|---|---|---|---|---|---|---|---|
| K23 | -- | 16.8 | 20.9 | 46.3 | 57.3 | 63.7 | 61.7 | 64.8 | 71.4 | 75.0 |
| K62 | 24.8 | -- | 11.5 | 30.1 | 53.4 | 53.2 | 53.1 | 55.4 | 62.8 | 66.5 |
| K68 | 22.1 | 11.5 | -- | 27.9 | 42.2 | 43.6 | 42.8 | 45.6 | 53.4 | 58.1 |
| K206 | 51.6 | 32.1 | 27.9 | -- | 50.3 | 35.3 | 39.6 | 39.2 | 47.5 | 51.2 |
| K537 | 62.5 | 56.2 | 44.5 | 50.7 | -- | 29.7 | 26.8 | 31.6 | 40.4 | 49.9 |
| K854 | 67.0 | 53.2 | 43.6 | 35.3 | 31.0 | -- | 8.7 | 7.3 | 19.0 | 28.0 |
| K857 | 63.3 | 53.1 | 42.8 | 39.6 | 26.8 | 12.2 | -- | 5.2 | 15.0 | 24.5 |
| K859 | 80.7 | 65.4 | 55.3 | 45.1 | 34.7 | 8.8 | 14.2 | -- | 12.0 | 21.4 |
| K867 | 86.2 | 75.0 | 64.4 | 56.9 | 43.2 | 24.4 | 20.9 | 18.8 | -- | 10.3 |
| K873 | 91.4 | 77.5 | 70.7 | 52.9 | 55.5 | 29.9 | 35.0 | 25.6 | 18.1 | -- |

PDB 3W5K

| | K23 | K62 | K68 | K206 | K537 | K854 | K857 | K859 | K867 | K873 |
|---|---|---|---|---|---|---|---|---|---|---|
| K23 | -- | 16.1 | 20.6 | 46.2 | 59.5 | 72.9 | 70.4 | 74.5 | 81.9 | 86.3 |
| K62 | 25.3 | -- | 11.7 | 30.7 | 56.4 | 66.0 | 65.4 | 69.0 | 77.8 | 82.8 |
| K68 | 25.2 | 12.9 | -- | 28.2 | 45.0 | 55.6 | 54.6 | 58.6 | 67.6 | 73.3 |
| K206 | 57.5 | 38.4 | 31.1 | -- | 53.7 | 52.5 | 56.1 | 58.2 | 69.1 | 75.1 |
| K537 | 66.3 | 60.0 | 47.4 | 56.1 | -- | 38.0 | 34.5 | 39.6 | 48.2 | 57.4 |
| K854 | 82.4 | 72.9 | 61.2 | 56.0 | 38.7 | -- | 8.9 | 7.5 | 19.1 | 27.7 |
| K857 | 73.8 | 71.9 | 59.0 | 57.2 | 34.5 | 12.3 | -- | 5.5 | 15.4 | 24.6 |
| K859 | 86.5 | 80.7 | 67.8 | 64.3 | 43.9 | 7.5 | 13.9 | -- | 12.0 | 21.1 |
| K867 | 95.0 | 93.0 | 80.1 | 77.8 | 49.8 | 21.1 | 19.1 | 16.6 | -- | 10.0 |
| K873 | 100.3 | 97.5 | 91.5 | 81.3 | 63.4 | 32.5 | 32.6 | 28.0 | 14.5 | -- |

**c**

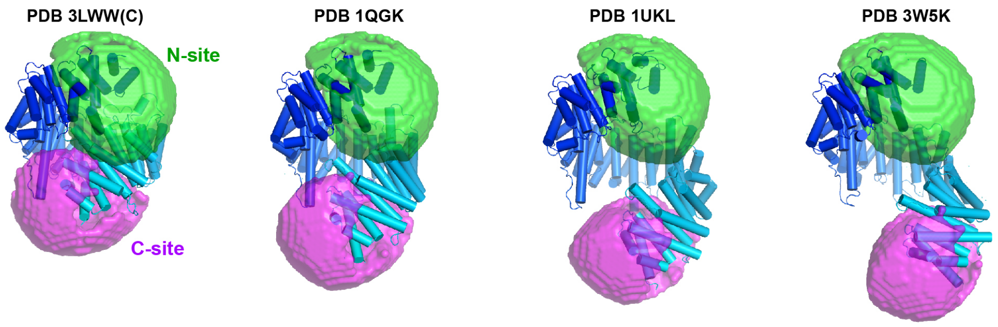


**Figure S3. Crosslinking-MS distance constraints localize two Rev-binding regions on Impβ. a.** Spheres of radius 35 Å centered on the Cα atoms of Group-1 residues Lys23 and Lys873 show that BS3 molecules bound to these two lysines cannot crosslink to the same Rev Lys position regardless of the Impβ conformation. The four conformations shown are those of Impβ bound to the following cargos: the IBB domain of Snurportin1 (PDB 3LWW chain C), the IBB domain of Importin α (PDB 1QGK), the SREBP-2 complex (PDB 1UKL) and Snail1 (PDB 3W5K). **b.** Distances between pairs of Impβ Group-1 Lys residues. The upper and lower triangles show distances for the conformations of Impβ bound to the IBB domain of Importin α (PDB 1QGK) and the SREBP-2 complex (PDB 1UKL), respectively. Solvent-accessible surface distances (SASDs) over 70 Å are outlined in dark blue. Distances were calculated using the Jwalk webserver[98]. **c.** Localization of the Cα atom of Rev Lys20. Green and magenta volumes show the N- and C-terminal regions of space within crosslinking distance of Group-1 residues in either HEAT repeats 1 and 2 (K23, K62, K68) or repeat 19 (K854, K857, K859, K867 and K873), respectively, defined by the intersection of 35 Å spheres centered on the Cα atoms of these residues.

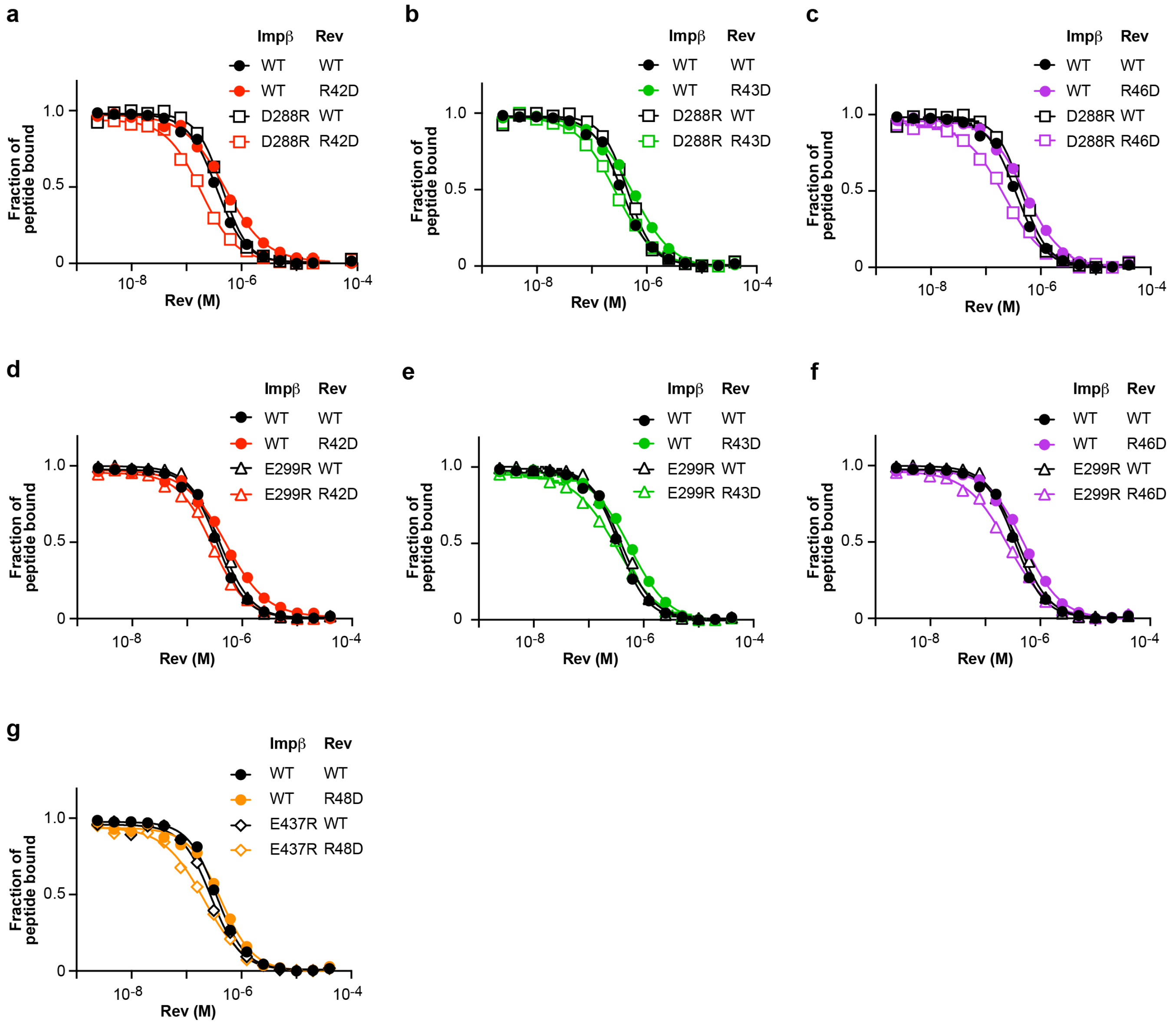


**Figure S4. Compensatory effects between charge-reversal mutants of Impβ and Rev.** Panels **a-g** show representative individual FP inhibition assays performed with WT Rev (black curves and symbols) or the indicated Rev mutant (colored curves and symbols) together with WT Impβ (circles) or the indicated Impβ mutant (squares, triangles or diamonds).

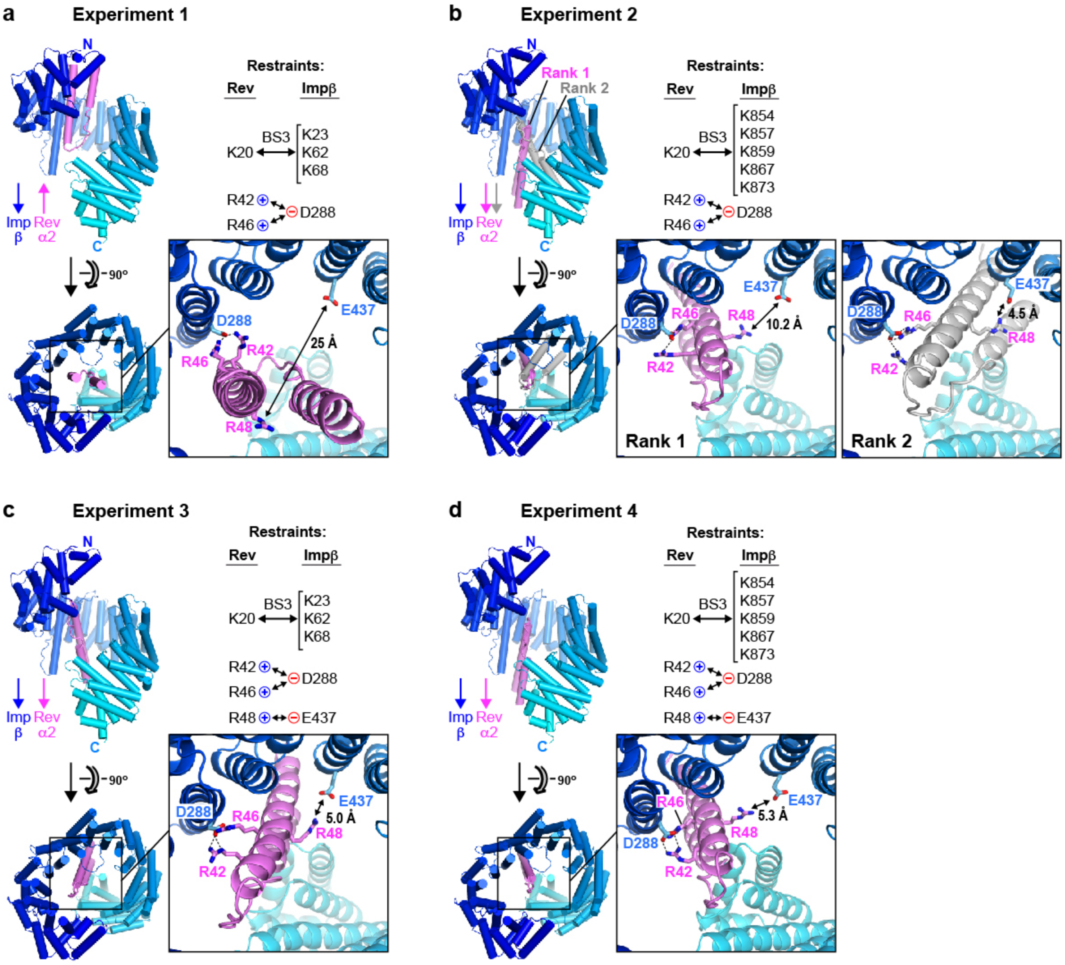


**e**

| Experiment | BS3 crosslinks: N-site | BS3 crosslinks: C-site | Electrostatic restraints (Rev: R42 / Impβ: D288) | Electrostatic restraints (Rev: R46 / Impβ: D288) | Electrostatic restraints (Rev: R48 / Impβ: E437) | Outcome | |
|---|---|---|---|---|---|---|---|
| 1 | + | - | + | + | - | **Expt. 3 worse than Expt. 1:** - fewer clustered solutions (84 vs 143) - greater diversity of configurations - Rev R48 and Impβ E437 are far apart | **Electrostatic restraints are more consistent with C-site than with N-site** |
| 3 | + | - | + | + | + | | |
| 2 | - | + | + | + | - | **Expt. 4 better than Expt. 2:** - more clustered solutions (189 vs 177) - higher uniformity of configurations - Rev R48 and Impβ E437 are close | |
| 4 | - | + | + | + | + | | |

**Figure S5. Electrostatic interactions suggested by compensatory mutagenesis data are more likely to involve Rev bound at the C-site than at the N-site.** Panels **a-d** show the results of HADDOCK docking experiments 1-4, respectively, described in the text. In each case the lowest-energy structure (L.E.S.) for the top-ranking cluster of docking solutions is shown, with Impβ colored from blue to cyan from N- to C-terminus and Rev in violet. In panel **b** the L.E.S. from the second ranked cluster is also illustrated, with Rev shown in gray. The pink and blue arrows indicate the orientation (from N- to C-terminus) of Rev helix α2 and the Impβ superhelical axis, respectively. BS3 crosslinking and electrostatic interaction restraints included in each experiment are summarized (see **Table S2a** for details). **e.** Evidence supporting the conclusion that electrostatic interactions deduced from our compensatory mutagenesis data are more compatible with the C-site of Impβ than with the N-site. Experiments 3 and 4 are identical to experiments 1 and 2, respectively, except that the $Glu437^{Imp\beta}$:$Arg48^{Rev}$ interaction was included as an additional distance restraint. Experiments 2 and 4 yielded similar results (**Table S2b**), except that experiment 4 resulted in a greater number of clustered solutions (189 versus 177) that were more uniform in configuration (compare **Figure S6d** with **S6b**) and yielded a top-ranked solution in which residues $Glu437^{Imp\beta}$ and $Arg48^{Rev}$ were closer together (compare panel **d** versus **b**). In contrast, experiment 3 yielded a different outcome compared to experiment 1 (**Table S2b**), resulting in fewer clustered solutions (84 versus 143) that were more diverse in configuration (compare **Figure S6c** with **S6a**). Indeed, several clusters of solutions (including the top-ranked) positioned Rev next to HEAT repeats 7-19 with helix α2 parallel to the Impβ superhelix, i.e., closely resembling the C-site solution for experiment 4 (panels **a,c** and **d**), even though the BS3 crosslinking restraints used were those associated with the N-site. Moreover, all the solutions that placed Rev in the N-site (clusters ranked $2^{nd}$ and $4^{th}$) positioned residues $Glu437^{Imp\beta}$ and $Arg48^{Rev}$ far apart. These findings suggest that the set of electrostatic interactions used as distance restraints in experiments 3 and 4 are more compatible with the C-site than with the N-site.

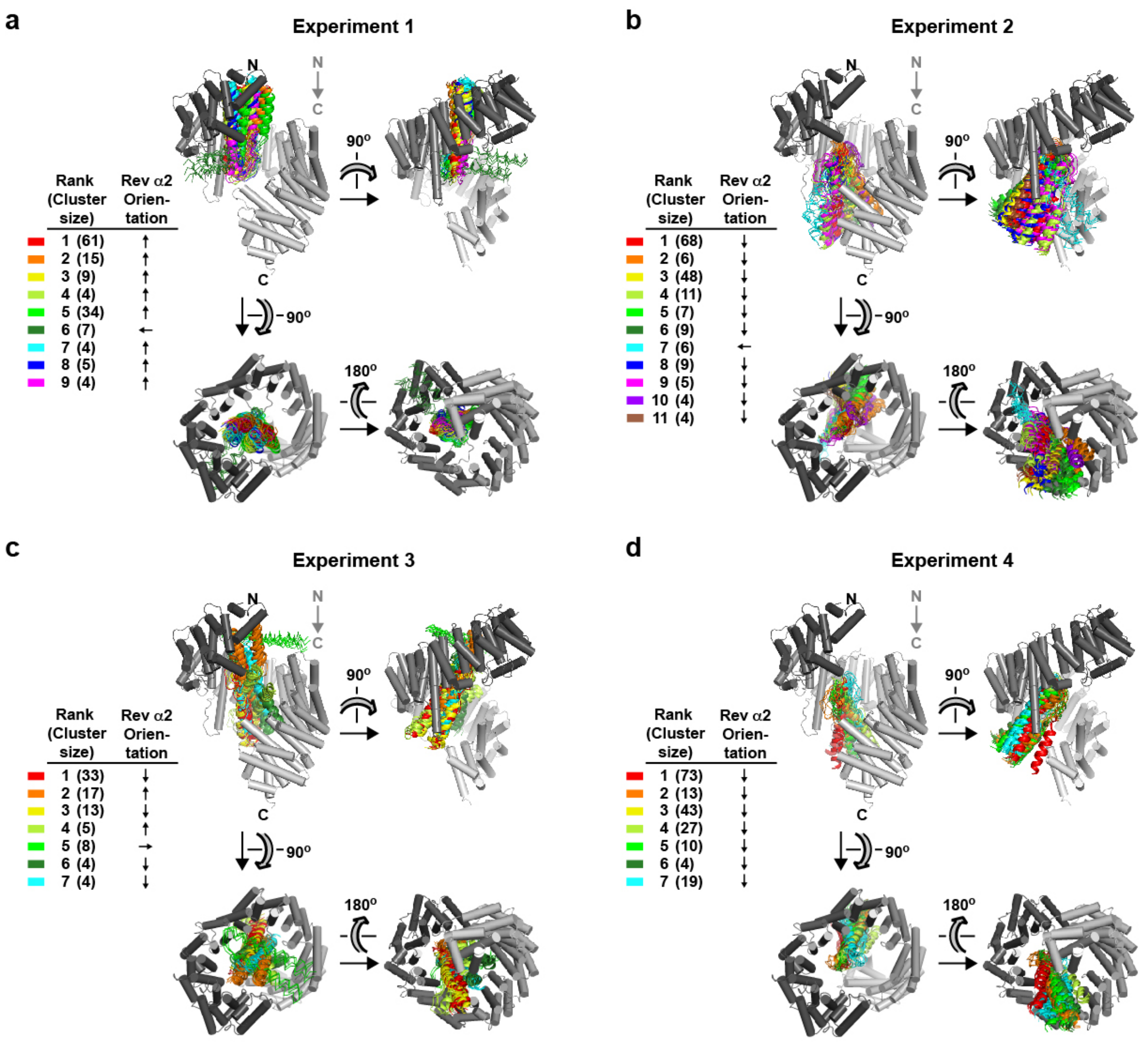


**Figure S6. Results of docking experiments with program HADDOCK.** Panels **a**-**d** show the results for Experiments 1-4, respectively. For each docking experiment, the 200 lowest-energy binding configurations were grouped into clusters according to structural similarity. Clusters were then ranked according to the average energy of the four lowest-energy structures in each cluster. The four lowest energy structures are shown for each cluster. All members of each cluster are colored according to the color scheme shown at the left. The downward gray arrow indicates the direction of the Impβ superhelical axis. For each cluster the approximate binding orientation of Rev is designated by indicating the orientation of Rev helix α2 as roughly antiparallel (↑), parallel (↓) or perpendicular (← or →) relative to the Impβ superhelical axis. Rev monomers with parallel or antiparallel orientations are shown as ribbon diagrams, those with perpendicular orientations are shown as Cα line tracings. Whereas the inclusion of the additional distance restraint between $Glu437^{Imp\beta}$ and $Arg48^{Rev}$ in Experiment 3 results in a greater diversity of cluster configurations relative to Experiment 1, its inclusion in Experiment 4 results in more uniform clusters relative to Experiment 2.

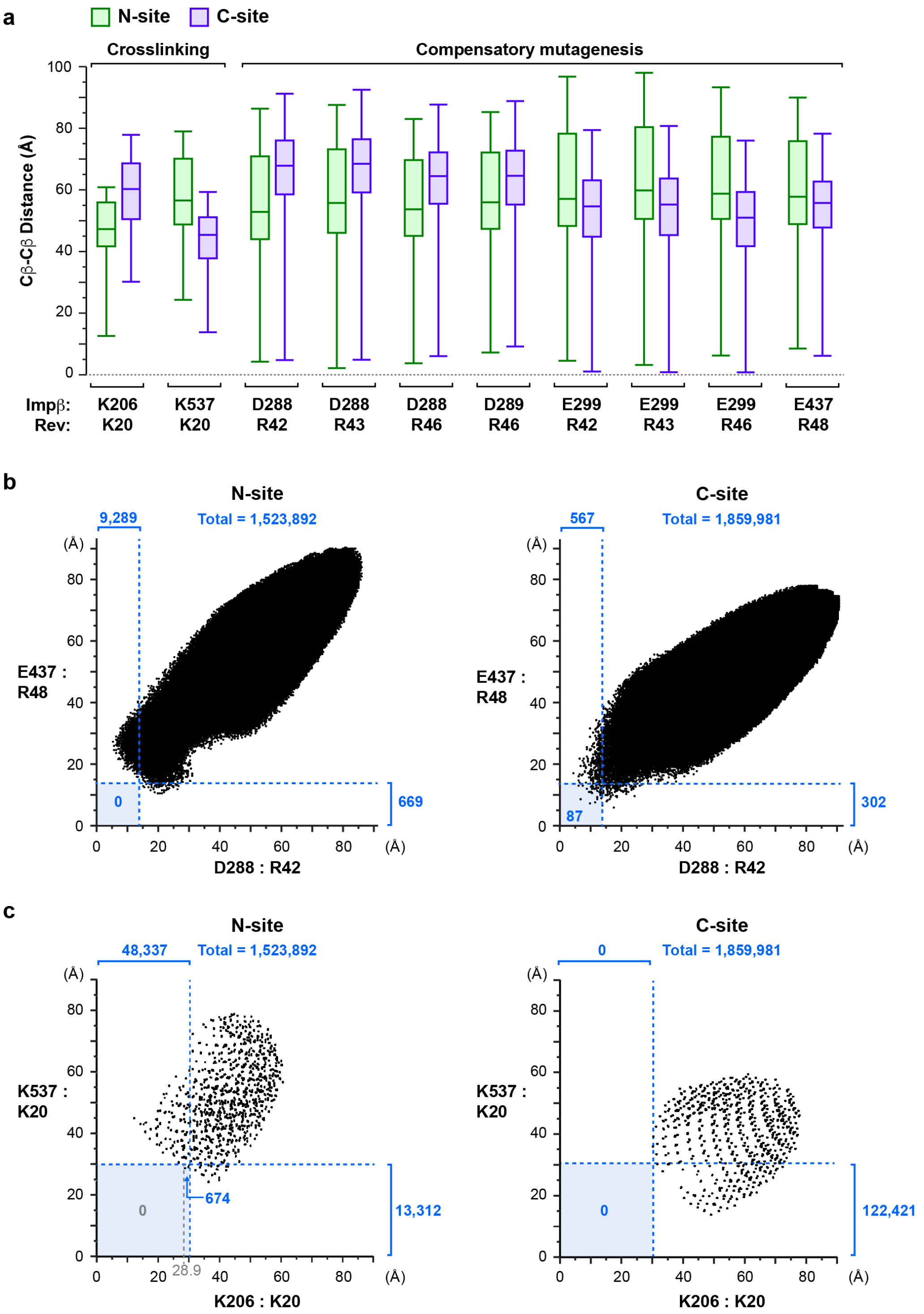


**Figure S7. Analysis of distances between Impβ and Rev residues in rigid-body docking simulations. a.** Boxplots of Cβ-Cβ distances between the indicated Impβ and Rev residues for docking simulations involving the N-site (green) or C-site (purple). **b.** Scatter plot of docking configurations showing the distribution of Cβ-Cβ distances between residues D288$^{Imp\beta}$ and Arg42$^{Rev}$ (horizontal ordinate) and between residues E437$^{Imp\beta}$:Arg48$^{Rev}$ (vertical ordinate) for docking simulations involving the N-site (left panel) or C-site (right panel). The number of configurations satisfying the indicated distance cutoffs are indicated in blue. **c.** Scatter plot of docking configurations showing the distribution of Cβ-Cβ distances between residues Lys206$^{Imp\beta}$ and Lys20$^{Rev}$ (horizontal ordinate) and between residues Lys537$^{Imp\beta}$:Lys20$^{Rev}$ (vertical ordinate) for docking simulations involving the N-site (left panel) or C-site (right panel). The number of configurations satisfying the distance cutoffs (indicated by dashed blue lines) are shown in blue. Because the Lys20$^{Rev}$ Cβ atom was centered on each grid position prior to applying the rotational search, each point in the plot may correspond to several different (up to 4056) Rev orientations.

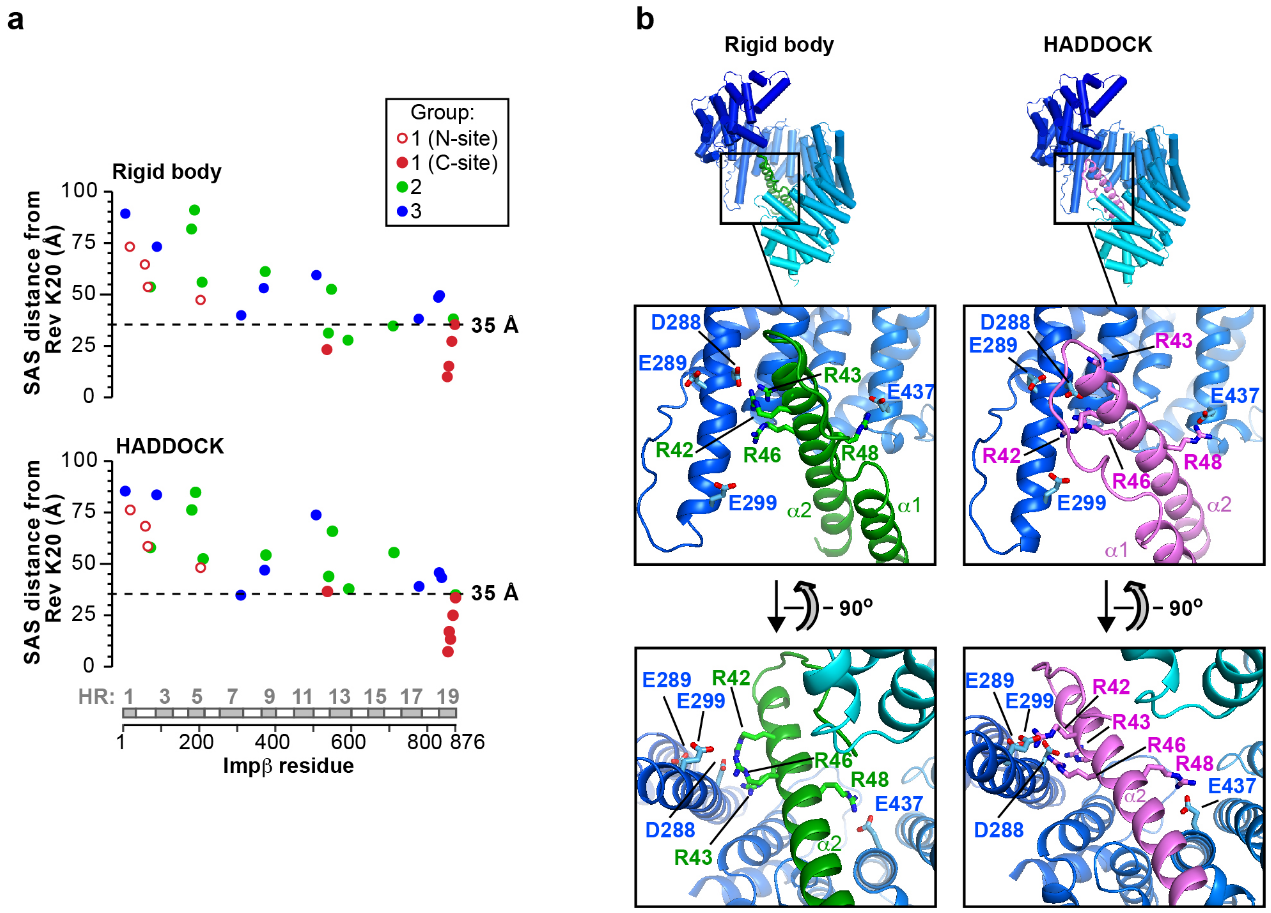


**Figure S8. Top-ranking configurations from docking experiments. a**. Plot of solvent-accessible surface (SAS) distances calculated for the top-ranking configuration obtained by rigid-body sampling (*top*) or HADDOCK (*bottom*) analysis. SAS distances were calculated between Rev residue Lys20 and Impβ lysine residues belonging to Group 1 (red), Group 2 (green) or Group 3 (blue), as defined in Figure 6b. Group 1 lysines associated with the N- or C-site of Impβ are indicated by open and closed red circles, respectively. SAS distances were calculated using the Jwalk server[98]. **b.** Orthogonal views of the top-ranking configuration obtained by rigid-body sampling (*left)* or HADDOCK (*right*) analysis. Acidic Impβ residues and basic Rev residues hypothesized to be in close proximity on the basis of compensatory charge-reversal mutation data are shown in stick representation. In the bottom panels Rev helix α1 is omitted for clarity.

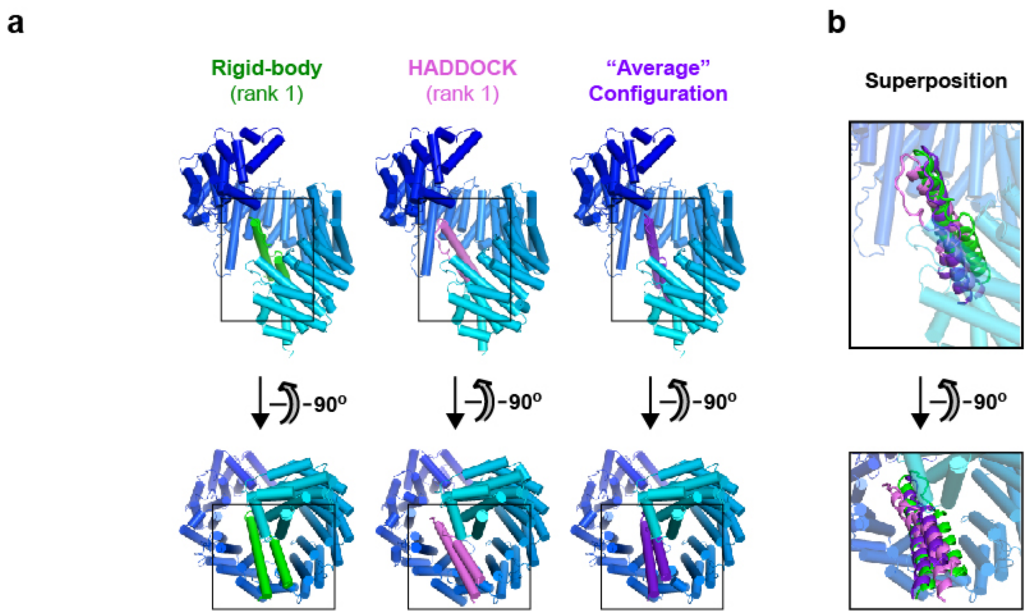


| Difference versus: | | relative rotation | |
|---|---|---|---|
| Rigid-body (rank 1) | - | **45.9°** | **25.9°** |
| HADDOCK (rank 1) | **3.0 Å** | - | **27.3°** |
| Average Configuration | **3.0 Å** | **1.0 Å** | - |
| | relative shift | | |

**Figure S9. Comparison of docking solutions. a.** *Top.* Structures of the top-ranking solution from the rigid-body docking and HADDOCK experiments are shown as well as the "average" configuration of all solutions obtained by the two docking methods. *Bottom.* Table summarizing the relative shift in the centroid position of Rev and its relative rotation (polar angle κ) in pairwise comparisons of the three structures, calculated using program Lsqkab from the CCP4 suite[126].

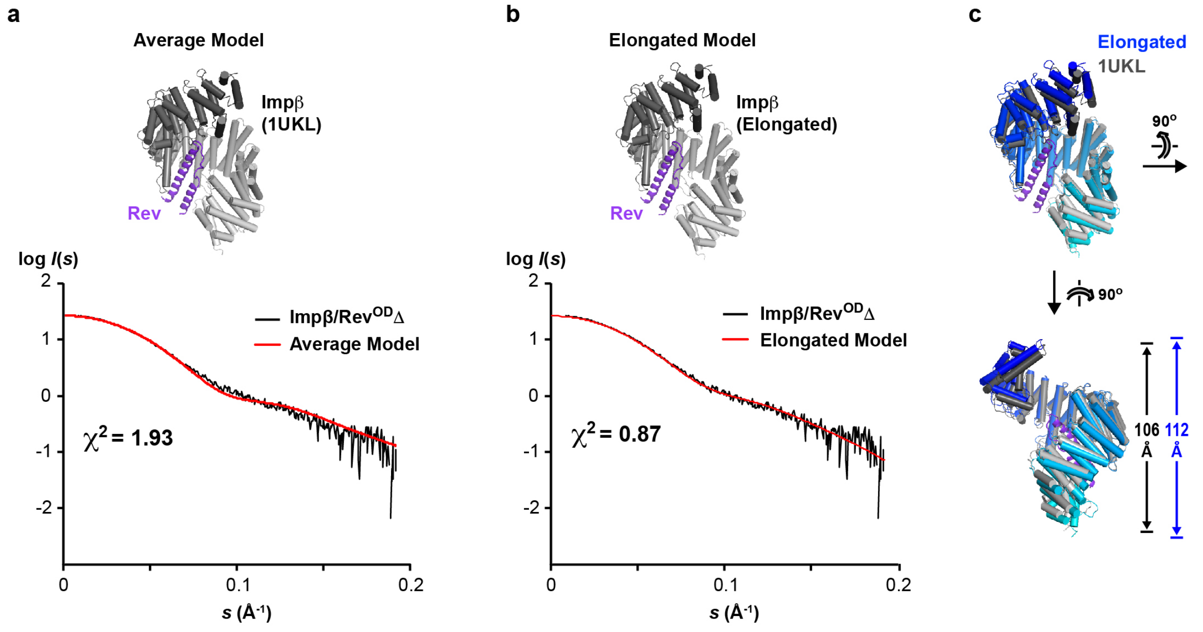


**Figure S10. The structural model of Rev bound to the C-site of Impβ complex agrees with SAXS data on the Impβ/Rev complex. a.** Scattering data from the Impβ/Rev$^{OD}\Delta$ complex compared to the scattering profile calculated from the "average" docking configuration of Rev bound to the Impβ C-site. **b.** A slightly more elongated Impβ conformation in the docking model yields an improved agreement between the observed and calculated scattering profiles. The elongated conformation was generated by normal mode perturbation of PDB 1UKL along the first (lowest-frequency) vibrational mode using the ElNemo webserver[125]. **c.** Comparison of the initial conformation of PDB 1UKL with the elongated conformation obtained by normal mode perturbation. The elongated conformation is more extended by ~6 Å along the HEAT-repeat superhelical axis.

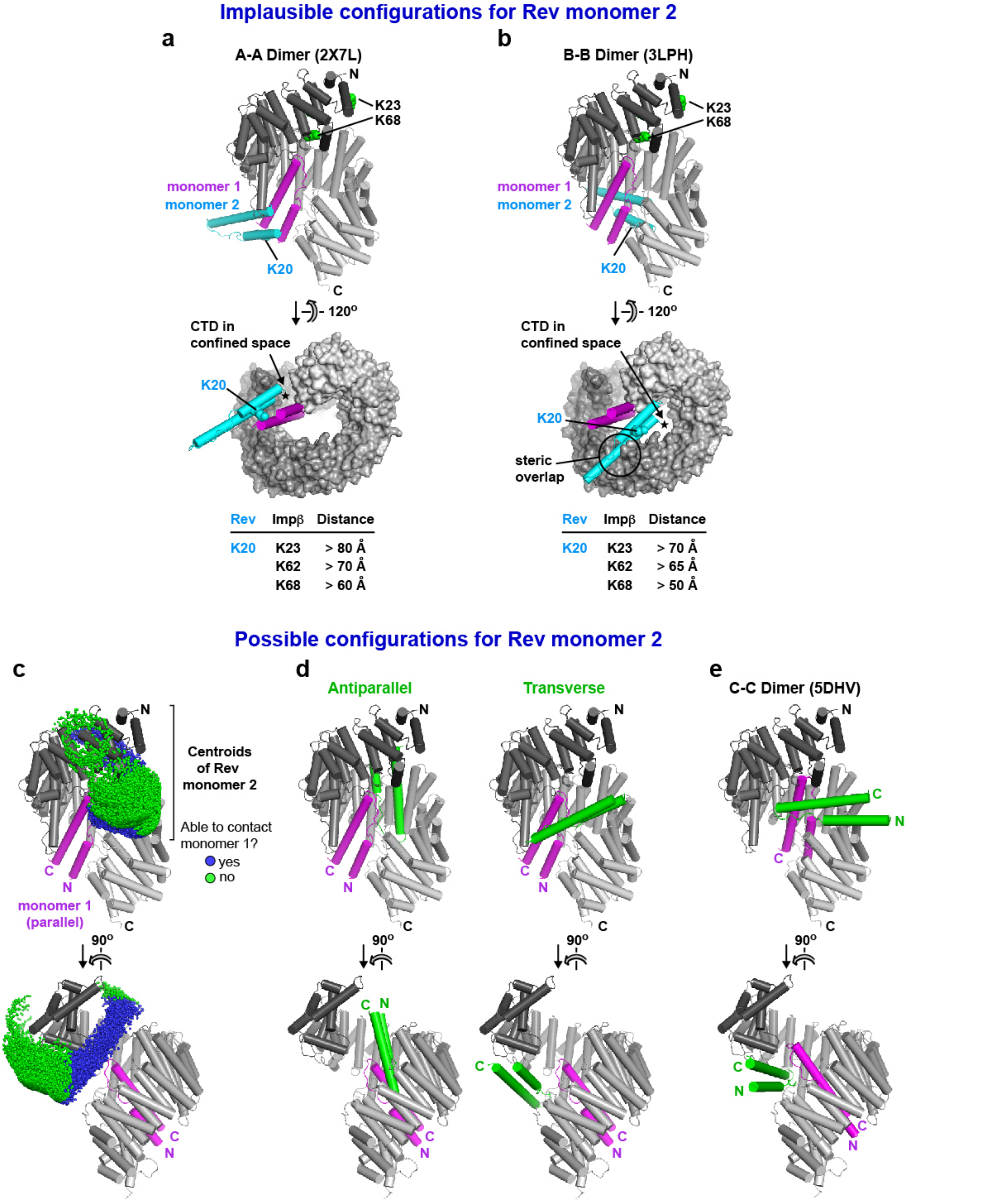


**Figure S11. Configurations for the binding of Rev monomer 2. a,b.** Unlikely configurations in which Rev monomer 2 (cyan) interacts with Rev monomer 1 (magenta) via an A-A (head-to-head) or B-B (tail-to-tail) interaction. Predicted distances between residue Lys20 on Rev monomer 2 and Impβ residues Lys23, Lys62 and Lys68 are roughly twice the BS3 crosslinking distance constraint. **c.** Results of the rigid body docking analysis showing the centroids of Rev monomer 2 that are consistent with BS3 crosslinking constraints associated with the N-site and sterically compatible with Impβ bound to Rev monomer 1 via the C-site. Centroids of Rev molecules within and beyond contact distance of Rev monomer 1 are indicated by blue and green spheres, respectively. **d.** Two representative orientations of Rev monomer 2 in which the N-and C-termini of the Rev helical hairpin are near the N-terminal end of Impβ (N-ward) or are located midway between the N- and C-terminal ends (Sideward). **e.** Hypothetical configuration showing that the binding of Rev to the N- and C-sites of Impβ is compatible with a C-C interaction between the two Rev monomers as observed in PDB 5DHV [67].

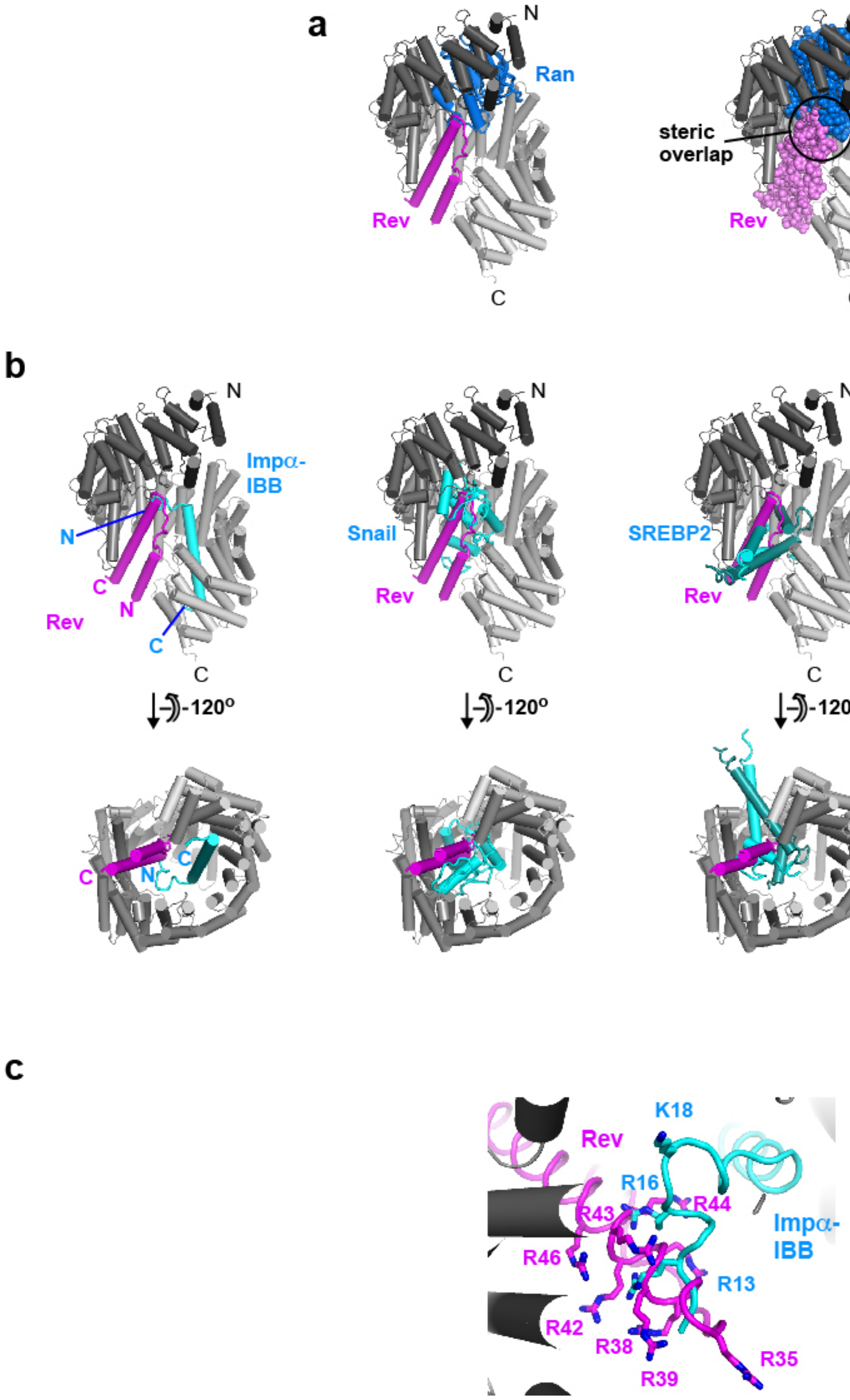


**Figure S12. Comparison of Impβ/Rev structural model with RanGTP- and cargo-bound complexes of Impβ. a.** Rev bound to the Impβ C-site is predicted to overlap sterically with RanGTP. **b.** Comparison with structures of cargo-bound Impβ complexes. **c.** The Rev ARM motif is predicted to localize to the same volume as the N-terminal moiety of the Impα IBB domain.
